# From gas to sugar: Trehalose production in *Cupriavidus necator* from CO_2_ and hydrogen gas

**DOI:** 10.1101/2020.06.05.136564

**Authors:** Hannes Löwe, Marleen Beentjes, Katharina Pflüger-Grau, Andreas Kremling

## Abstract

The paradigm of a fossil based, non-renewable economy will have to change in the future due to environmental concerns and the inevitable depletion of resources. Therefore, the way we produce and consume chemicals has to be rethought: The bio-economy offers such a concept for the sustainable production of commodity chemicals using waste streams or renewable electricity and CO_2_. Residual biomass or organic wastes can be gasified to energy rich mixtures that in turn can be used for synthesis gas fermentation.

Within this scope, we present a new process for the production of trehalose from gaseous substrates with the hydrogen-oxidizing bacterium *Cupriavidus necator* H16. We first show that *C. necator* is a natural producer of trehalose, accumulating up to 3.6% of its cell dry weight as trehalose when stressed with 150 mM sodium chloride. Bioinformatic investigations revealed a so far unknown mode of trehalose and glycogen metabolism in this organism. Next, we evaluated different concepts for the secretion of trehalose and found that expression of the sugar efflux transporter A (*setA*) from *Escherichia coli* was able to lead to a trehalose-leaky phenotype. Finally, we characterized the strain under autotrophic conditions using a H_2_/CO_2_/O_2_-mixture and other substrates. Even without overexpressing trehalose synthesis genes, titers of 0.47 g/L and yields of around 10% were reached, which shows the great potential of this process.

Taken together, this process represents a new way to produce sugars with a higher areal efficiency than photosynthesis by crop plants. With further metabolic engineering, we anticipate an application of this technology for the renewable production of trehalose and other sugars, as well as for the synthesis of ^13^C-labeled sugars.

**Graphical abstract:** 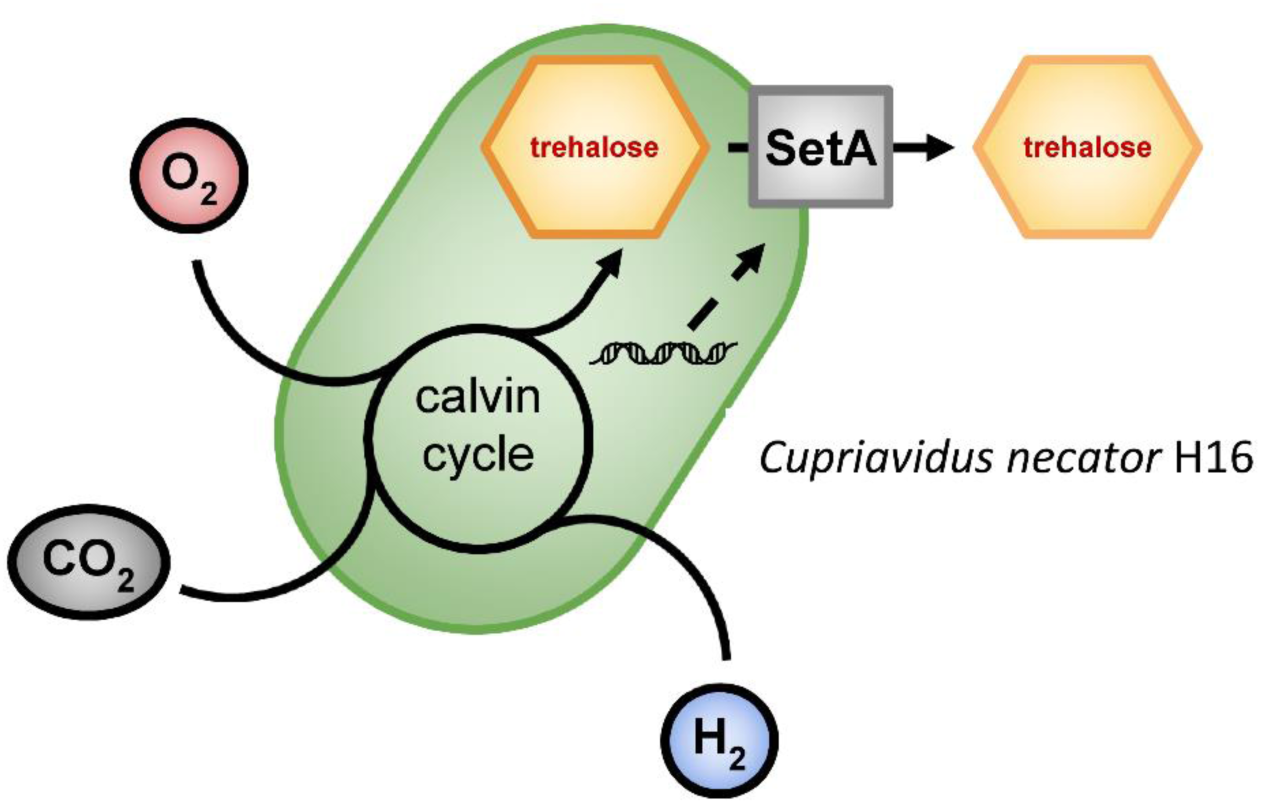

## Introduction

Transforming our fossil resource based economy into a sustainable one, as currently planned by many industrial countries (European Commission, 2020), will come along with massive changes in the way energy and chemicals are produced and used. This holds great potential for biotechnological processes since nature supplies us with an easy and readily available renewable resource in form of crop plants. Although these plants provide sugars and fats that can be directly fed to biotechnological processes, they also have downsides, mainly stemming from their large areal demand: i) the amount of light that is fixed into chemical energy of the desired substrate is rather low (2.4-3.7% (Zhu et al., 2008) compared to > 14% with photovoltaics coupled with electrolysis (Rothschild & Dotan, 2017)), ii) there is a conflict of the production of food with substrates for biotechnology because both use the same agricultural area, iii) industrial cultivation of crops poses threats to our ecosystems in form of water consumption, erosion, pesticides (Woodhouse, 2010) and many more.

One strategy to mitigate these problems is to use alternative renewable resources: surplus glycerol has been widely used as a substrate for biotechnological processes and residual biomass can be hydrolyzed into its sugar monomers or directly gasified to produce energy-rich synthesis gas. This energy-rich gas can then in turn be converted to methanol, which is the feed-stock for the so-called “methanol-economy”, or directly fed to microbes for synthesis gas fermentation. These concepts are under research by a great number of research groups and have also partially been commercialized (Sun et al., 2019). Energy rich gas in form of hydrogen can also be produced in a sustainable way by coupling photovoltaics to electrolysis that have a substantially higher areal efficiency than crops. Hydrogen gas can be mixed with CO_2_ and used as synthesis gas. Alternatively, it can be used to increase the hydrogen content of synthesis gas derived from gasified biomass or waste, (Clausen et al., 2010). CO_2_ can also be hydrogenated to formic acid that is the substrate for the “formate based bio-economy” (Yishai et al., 2016). Alternatively, the direct electro-catalytic production of formic acid is investigated by some research groups (S. Lee et al., 2015). Synthesis gas, formate and methanol can be used by acetogens or methanogens in anaerobic processes to produce highly reduced organic compounds like short-chained fatty acids and alcohols or methane with high yield and energy efficiency (Claassens et al., 2019). Other biomolecules, like sugar, that have a high demand of ATP for their synthesis are, however, not suitable as products for these organisms because of thermodynamical reasons.

For this purpose, hydrogen-oxidizing bacteria are well suited as they can use synthesis gas and at same time are able to breath oxygen to generate ATP that is necessary to produce these energy rich compounds. In this study we show for the first time the production of the disaccharide trehalose from non-carbohydrate substrates with the hydrogen-oxidizing bacterium *Cupriavidus necator* H16, formerly known as *Ralstonia eutropha* H16, by genetic engineering. First, we elucidate the natural occurring trehalose metabolism in this bacterium by bioinformatics methods and experimentally show the accumulation of said sugar. Next, we engineer it for the secretion of trehalose, resembling the approach applied previously (Ducat et al., 2012) for secretion of sucrose from cyanobacteria. And finally, we show production of trehalose from a hydrogen-gas enriched gas mixture and other, non-carbohydrate substrates with the engineered *C. necator* strain.

## Results and Discussion

### Bioinformatic identification of trehalose and glycogen synthesis genes

Recently, autotrophic sugar production and secretion have been engineered and demonstrated in cyanobacteria by various studies (Ducat et al., 2012; Lin et al., 2020; Thiel et al., 2019). An impressive proportion of the fixed carbon can be directed to sugar synthesis and exported from the cell. As both cyanobacteria and *C. necator* are using the Calvin Cycle under autotrophic conditions for carbon fixation, we assumed that a similar concept might be applied to *C. necator*. Homologs of genes for the synthesis of the cyanobacterial osmolytes sucrose or glycosylglycerol are missing in *C. necator*, however. Instead, we found gene clusters for the synthesis of trehalose (Fig. 2) which is accumulated by some Gram-negative bacteria under conditions of stress like elevated salt concentrations (Burg & Ferraris, 2008).

**Figure 1:**
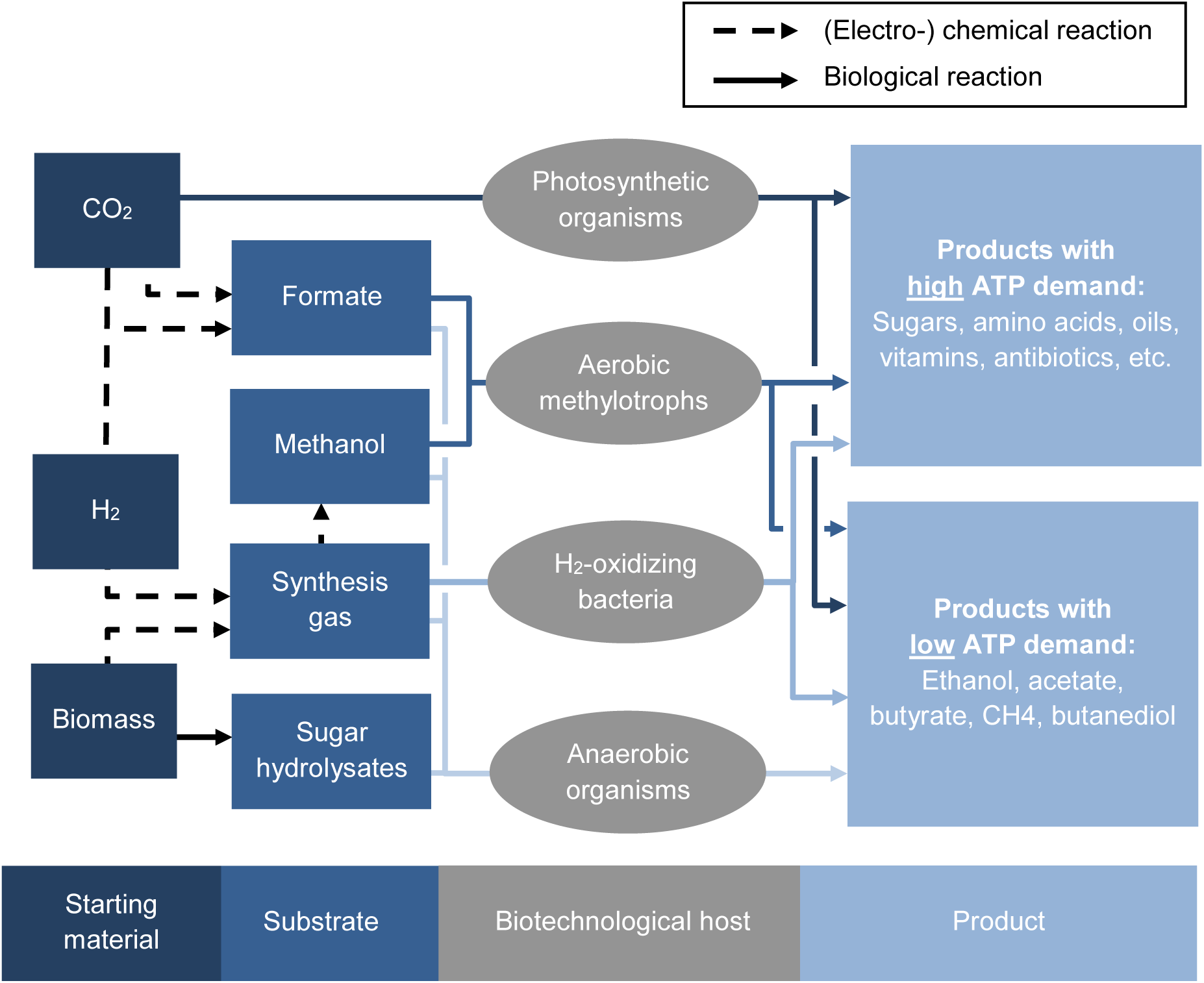
Existing and envisioned platforms for a renewable, future bioeconomy. Starting from different raw material, substrates for fermentation can be produced either with (electro-)chemical methods (dashed arrows) or with biological conversions (straight arrows). Formate will be the substrate for the so called “formate bioeconomy” and methanol for the “methanol bioeconomy”. Sugar hydrolysates can be regarded as a drop-in substrate for existing platform microorganisms. All aerobic platform organisms (the upper three groups) are able to potentially produce products with high ATP demand while anaerobic organisms like acetogens and methanogens are restricted to products with low ATP demand.

**Figure 2:**
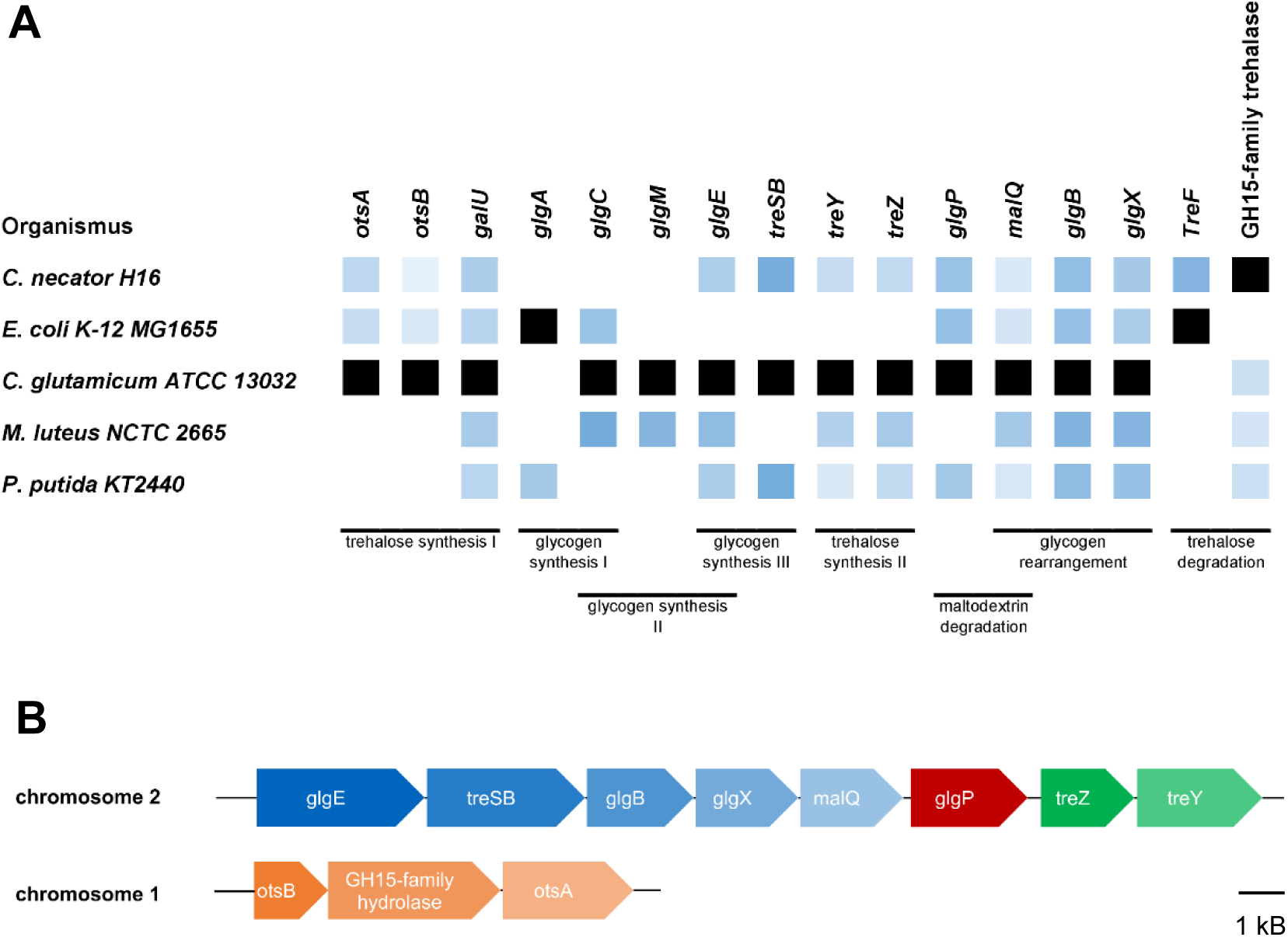
Homology search and annotation of genes involved in glycogen and trehalose metabolism. **A:** Co-occurance of homologous genes in different organisms in which glycogen and trehalose metabolism has been studied in the literature (Belda et al., 2016; Joseph et al., 2010; Kizawa et al., 1995; Wolf et al., 2003). Black color indicates the template that has been used for homology search. The shade of blue represents the homology score: Lighter shades of blue mean less homology; more saturated values mean closer homology. **B**: Arrangement of gene clusters in *C. necator*; all genes are organized in two clusters that are each possibly transcribed together since there are only small gaps (if any) between neighboring genes. Putative operons are indicated by different colors as the gaps between genes in different colors are large enough to contain a promoter. Length of blocks and gaps scale to the actual length of the genes.

There are three pathways for the synthesis of trehalose which all seem to be present in *C. necator*:

1. The *otsAB* pathway that is well described in *Escherichia coli*, is responsible for the production of trehalose from glucose-6-phosphate and UDP-glucose.
2. The *treYZ* pathway connects glycogen and trehalose metabolism and uses glycogen as a substrate for trehalose synthesis.
3. The trehalose synthase TreS(B) catalyzes the reversible conversion of maltose to trehalose, but is not suitable for trehalose accumulation since almost equal amounts of maltose would be co-accumulated. As maltose has a reactive aldehyde group in contrast to all known osmolytes, this is a rather unlikely way to produce trehalose. TreS instead seems to be involved in trehalose degradation/cycling (Ruhal et al., 2013).

All genes involved in trehalose synthesis are organized in two gene clusters (Fig. 2 B). The *otsAB* cluster is located on chromosome 1 and *treYZ, treSB* and the genes involved in glycogen synthesis are located in a gene cluster on chromosome 2. When comparing the genes for glycogen production in various organisms, it is interesting to note that *C. necator* does not contain any homolog of the *glgA* or *glgM* genes which are responsible for glycogen synthesis in all known organisms (Fig. 2 A). The only way to produce glycogen would be through the synthesis of trehalose by OtsAB, conversion to maltose-phosphate by TreSB and elongation of glycogen by GlgE (illustrated in Fig. 3). This pathway of glycogen synthesis has not been described in other organisms, but as trehalose and glycogen metabolism seem to be closely intertwined (Ruhal et al., 2013), it seems likely that the trehalose based synthesis of glycogen could be more wide-spread in bacteria. Since glycogen metabolism has never been experimentally addressed in *C. necator*, this needs further investigation.

**Figure 3:**
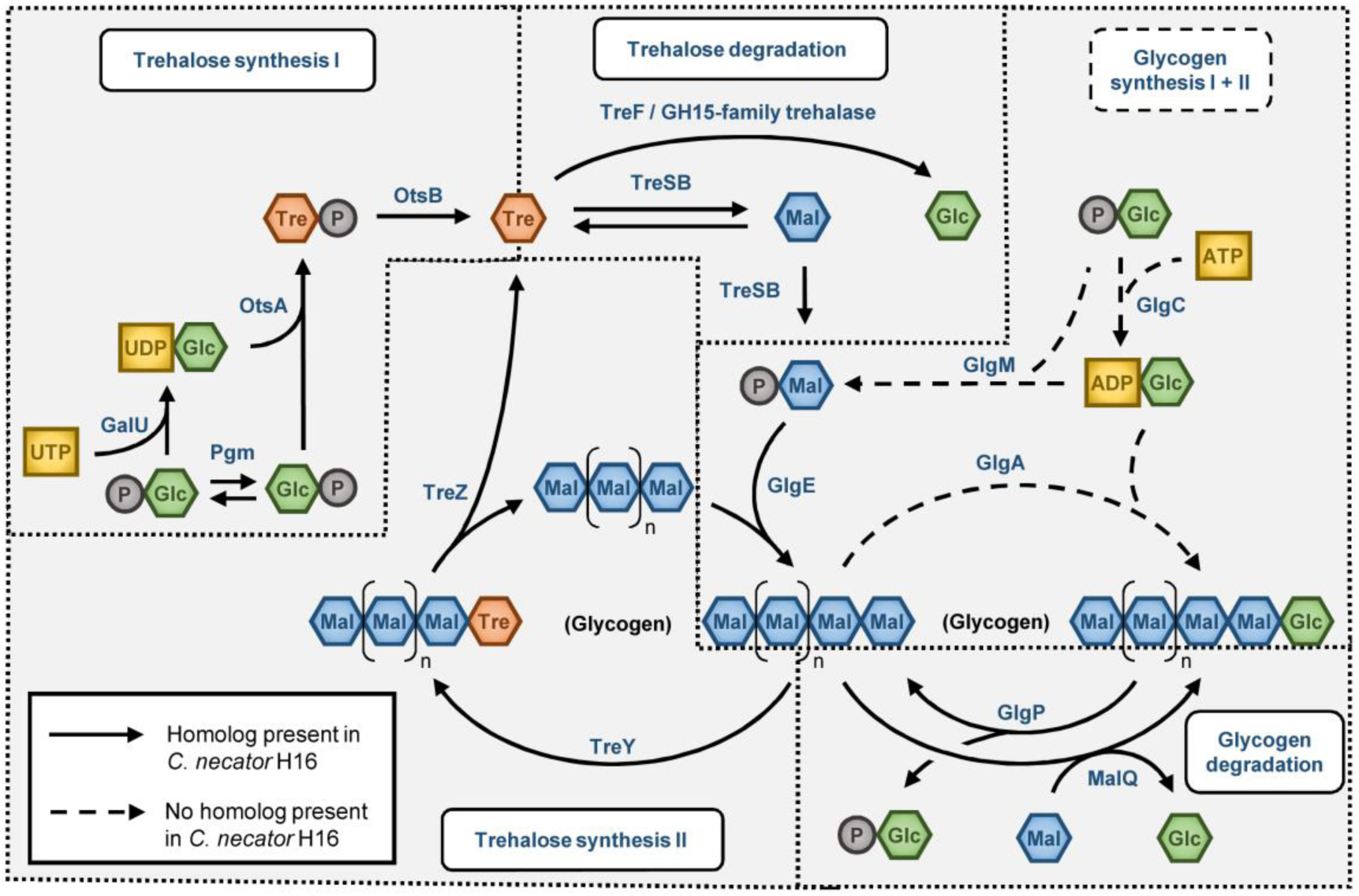
Pathways involved in trehalose and glycogen synthesis. Solid arrows indicate that a homolog for the protein catalyzing the reaction exists in *C. necator*, dashed arrows indicate the absence of an enzyme for the respective reaction. Note that all classical enzymes for glycogen synthesis are missing in *C. necator*.

Another striking feature of *C. necator* is the organization of its *ots* operon since it contains an additional gene in between the genes necessary for the synthesis of trehalose. This gene is auto-annotated as a glucoamylase of the glucoside hydrolase 15 family in the reference Genbank genome (accession number NC_008313). A homologous enzyme has been described as a Ca^2+^/Mg^2+^ and phosphate dependent trehalase in *Mycobacterium smegmatis* (Carroll et al., 2007). The gene of *C. necator* did not show any significant trehalase activity in cell lysates when expressed from a pSEVA228 expression plasmid in *E. coli* DH5α with or without Mg^2+^ and/or phosphate when compared to strains without the plasmid (data not shown). Additionally, as the simultaneous expression of the trehalose synthesis genes and a trehalase are counter-intuitive, we speculate that this gene has another, yet unknown function that could be uncovered in future studies.

The details of the bioinformatic analysis can be found in a table in the Supplementary Information of this study.

### *C. necator* H16 accumulates trehalose under conditions of salt stress

Next, we tested whether or not *C. necator* H16 is actually accumulating trehalose under laboratory growth conditions. In former publications, it was found that *E. coli* produces trehalose when cells are stressed by increased osmotic pressure (Burg & Ferraris, 2008). To test whether this is also true for *C. necator* H16, we performed experiments in shakeflasks with 150 mM NaCl or without salt. As a control we also checked the influence of 3-methylbenzoate (3-MB) on trehalose synthesis, since it is one of the preferred inducers for the XylS/Pm expression system that we later on used for gene expression in *C. necator* for further genetic engineering.

In fact, cell lysates of cultures that were grown over-night and analyzed by HPLC with sugars and organic acids column, showed a peak in response to salt addition with a retention time equal to a trehalose standard. To confirm that the peak corresponded indeed to trehalose, the samples were treated with cell lysate of cells expressing the cytoplasmic trehalase TreF from *E. coli*. Cleavage with TreF resulted in the formation of glucose as the sole product, validating that the accumulated compound was trehalose.

Table 1 summarizes the optical densities, the trehalose concentrations and estimated trehalose mass fraction of the cells at the time of sampling one day after inoculation.

**Table 1:** Trehalose formation in wild type *C. necator* H16 cells with and without salt stress

| Condition | OD600 | Trehalose*, g L<sup>-1</sup> | Mass fraction** g g<sup>-1</sup> |
| --- | --- | --- | --- |
| 0 mM NaCl | 3.6 | 0.035 | 0.0055 |
| 0 mM NaCl + 0.1 mM 3-MB | 3.6 | 0.030 | 0.0048 |
| 150 mM NaCl | 2.7 | 0.167 | 0.0358 |

\* projected to total culture volume
\*\* estimated from a CDW/OD600-correlation of 0.363 g L<sup>-1</sup> (Grunwald et al., 2015)

### Engineering *C. necator* H16 for trehalose secretion

Having shown that *C. necator* is able to accumulate trehalose intracellularly, we set out to genetically modify the strain to secrete the disaccharide into the medium, following the approach that has been demonstrated in cyanobacteria in previous works (Ducat et al., 2012): there, the osmolyte sucrose that is naturally produced by many bacteria in response to salt stress was secreted by means of the sucrose permease CscB from *E. coli*. Originally, CscB has been characterized as a sucrose/H^+^-symporter (Peng et al., 2009; Sahintoth et al., 1995) that transports sucrose into the cell driven by the force of the membrane potential. A trehalose/H^+^-symporter has unfortunately never been described in the literature, neither has an antiporter or channel protein been found. In *Drosophila melanogaster*, the trehalose facilitator Tret1-1 has been characterized (Kanamori et al., 2010). Functional expression of eukaryotic membrane proteins has been proven difficult in bacteria in many cases (He et al., 2014; Schlegel et al., 2010). Therefore, this was not the ideal protein candidate. In order to find a way to secrete sugars in *C. necator*, we tested three different transport proteins (illustrated in Fig. 4):

**Figure 4:**
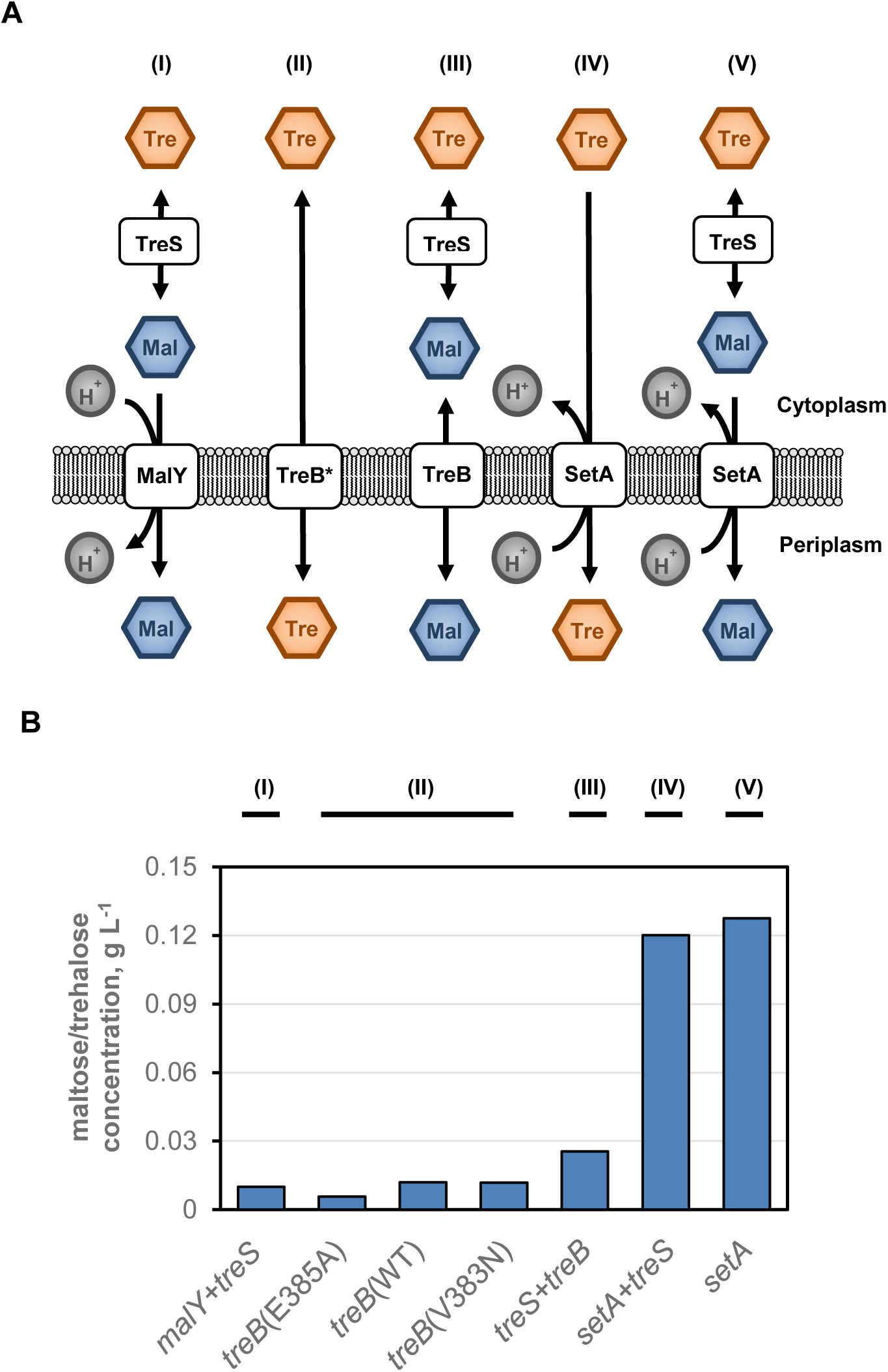
Screening of different transport proteins for the capability to secrete trehalose or maltose in *C. necator*. To secrete maltose, trehalose had to first be converted to maltose by the action of trehalose synthase TreS. All genes were expressed from pSEVA228-based plasmids, induced with 0.1 mM 3-MB. **A**) Schematic depiction of the tested strategies and assumed mode of action of the different transport enzymes. TreB* denotes the muted versions of TreB. **B**) Experimental characterization of the different genetic constructs that correspond to the strategies. Mutations of TreB are indicates in parentheses, the wildtype (WT) was used as a control.

1. Converting trehalose to maltose by trehalose synthase A from *Pseudomonas putida* KT2440 and secretion by the maltose/H^+^-symporter MalY that is found in *Neisseria meningitidis* (Derkaoui et al., 2016) (strategy (I) in Fig. 4). We used the *malY* gene from *Neisseria lactamica*, a close relative with safety class I. The two genes were combined in the bicistronic expression plasmid pSEVA228-malY-treS.
2. Engineering the trehalose PTS IIC subunit of *E. coli* DH5α (*treB* gene) for facilitated transport of trehalose (strategy (II) in Fig. 4). Other PTS transport subunits II have been shown to switch from phosphorylation to facilitated transport when amino acids in the highly conserved GITE-motif are mutated (Otte et al., 2003; Ruijter et al., 1992). In the TreB protein, the homologous sequence equaling to “GVTE” is spanning from amino acid position 382 to 385. We chose the two mutations E385A and V383N as they were successful for the mannitol PTS IIC transport subunit (Otte et al., 2003) and the glucose PTS subunit II *ptsG* in *E. coli* (Ruijter et al., 1992), respectively. The mutated versions of *treB* were constructed and ligated into the multiple cloning site of pSEVA228. As a control the wild type *treB* gene was also tested. On top of trehalose secretion, we also tested facilitated transport of maltose through TreB as reported by *Decker et al.* (Decker et al., 1999) (strategy (III) in Fig. 4). For this purpose, a bicistronic gene expression cassette with *treB* with *treS* was constructed.
3. Expression of the sugar efflux transporter gene setA from E. coli. The protein product has been shown to transport glucose and lactose (Liu, Miller, Willard, et al., 1999). In addition, several saccharides, among them maltose, were able to inhibit transport of lactose. It was speculated that these saccharides might also be substrates for SetA (Liu, Miller, Willard, et al., 1999) (strategy (IV) and (V) in Fig. 4). We cloned the *setA* gene from *E. coli* BL21 into pSEVA228 yielding pSEVA228-setA. Additionally, we combined *setA* with *treS* in the plasmid pSEVA228-setA-treS to check also for a possible secretion of maltose alone.

A complete list of tested expression plasmids can be found in Table 4. All plasmids were evaluated for their ability to confer sugar secretion to *C. necator* with 150 mM NaCl and induced with 3-MB using fructose as a carbon source. In Fig. 4B the trehalose in the supernatant after one day of cultivation is shown.

**Table 2:**
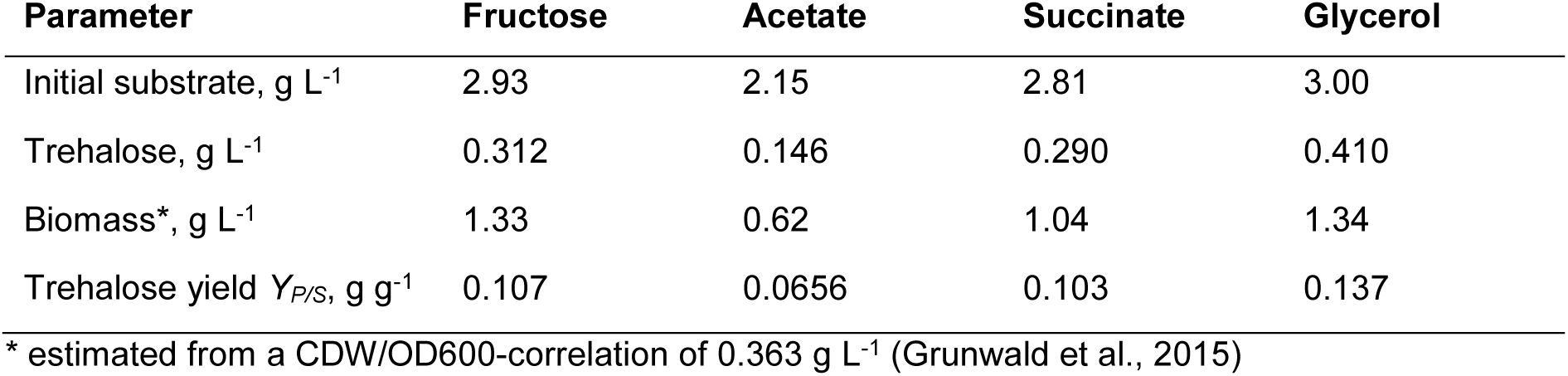
Product concentrations, cell dry weights and yields for cultivations with *C. necator*

| Parameter | Fructose | Acetate | Succinate | Glycerol |
| --- | --- | --- | --- | --- |
| Initial substrate, g L<sup>-1</sup> | 2.93 | 2.15 | 2.81 | 3.00 |
| Trehalose, g L<sup>-1</sup> | 0.312 | 0.146 | 0.290 | 0.410 |
| Biomass*, g L<sup>-1</sup> | 1.33 | 0.62 | 1.04 | 1.34 |
| Trehalose yield $Y_{P/S}$, g g<sup>-1</sup> | 0.107 | 0.0656 | 0.103 | 0.137 |

\* estimated from a CDW/OD600-correlation of 0.363 g L<sup>-1</sup> (Grunwald et al., 2015)

**Table 3:**
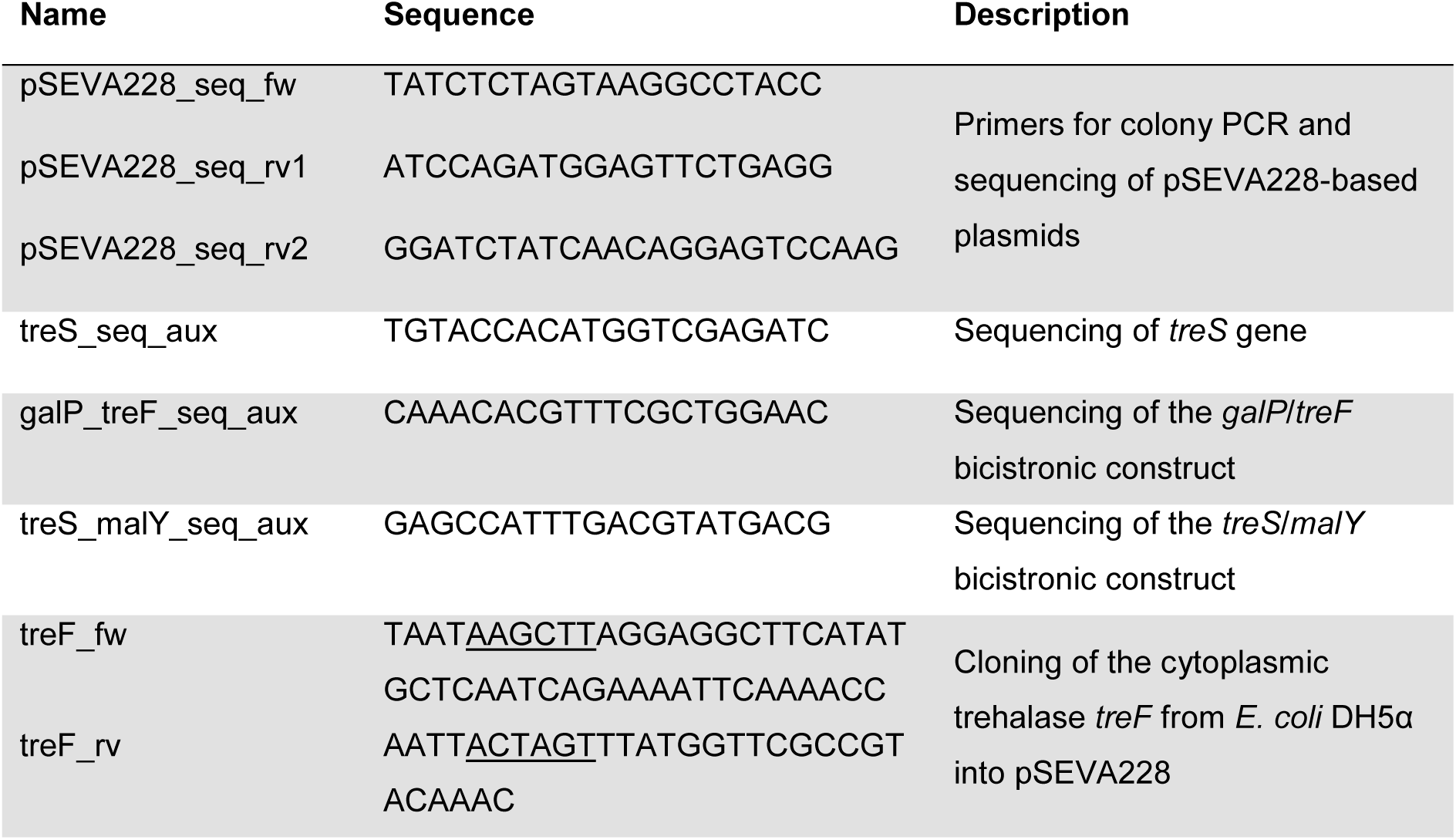
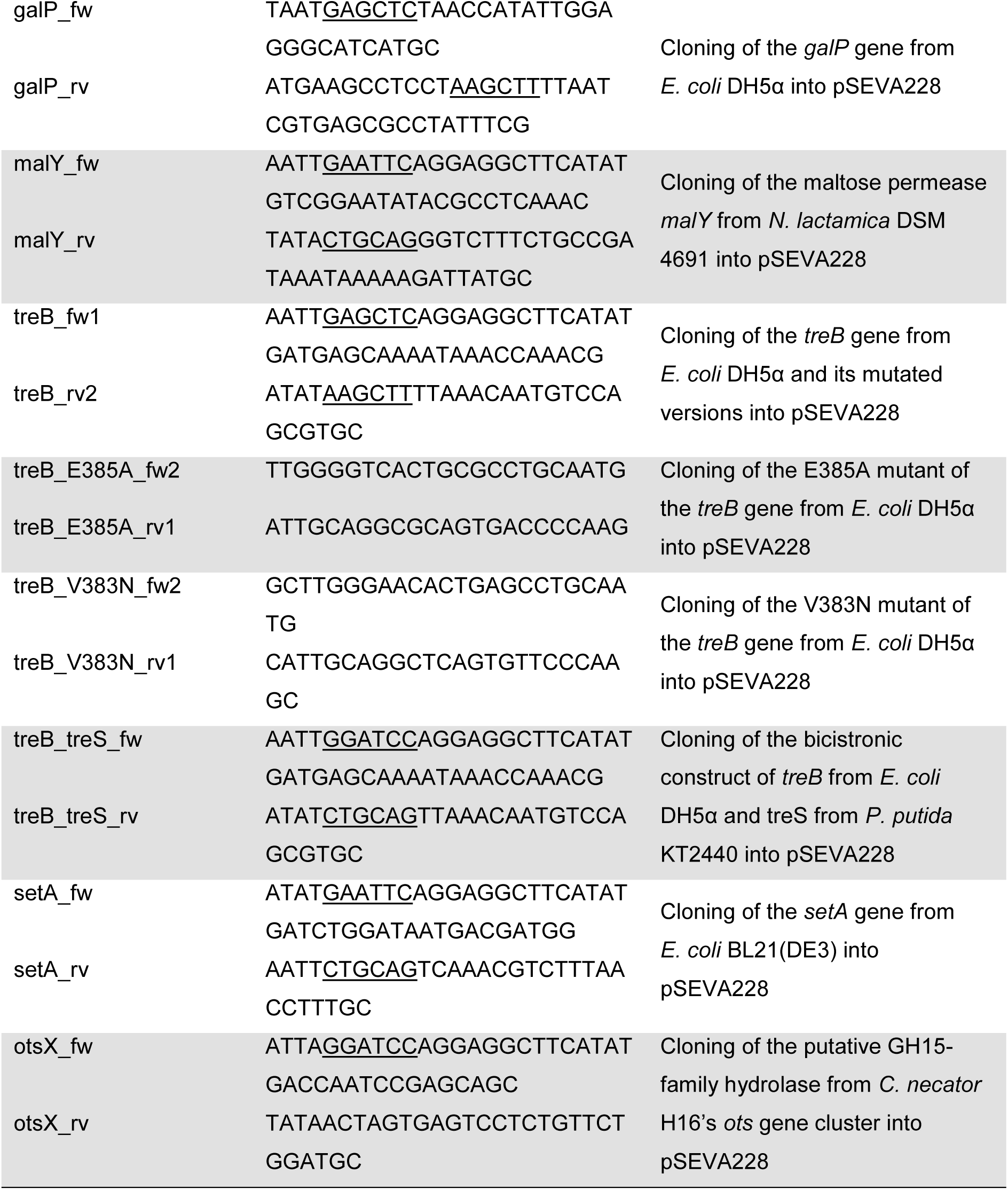
List of oligonucleotides used in this study. Restriction sites are underlined.

**Table 4:** List of all plasmids used in this study

| Plasmid name | Description | Source or reference |
| --- | --- | --- |
| pSEVA228 | Empty XylS/Pm expression plasmid with neomycin resistance and RK2 ori | (Martínez-García et al., 2015) |
| pRK600 | Helper plasmid for conjugative transfer of plasmids | (Kessler et al., 1992) |
| pSEVA228-treB | Expression plasmid for *treB* from *E. coli* DH5α | this study |
| pSEVA228-treB(V383N) | Expression plasmid for *treB* from *E. coli* DH5α, with the mutation V383N | this study |
| pSEVA228-treB(E385A) | Expression plasmid for *treB* from *E. coli* DH5α, with the mutation V383N | this study |
| pSEBA228-treB-treS | Expression plasmid for *treB* from *E. coli* DH5α, combined with a *treSA* from *P. putida* KT2440 in bicistronic construct | this study |
| pSEVA228-malY-treS | Expression plasmid for *malY* from *N. lactamica* DSM 4691, combined with a *treSA* from *P. putida* KT2440 in bicistronic construct | this study |
| pSEVA228-setA | Expression plasmid for *setA* from *E. coli* BL21(DE3) | this study |
| pSEVA228-setA-treS | Expression plasmid for *setA* from *E. coli* BL21(DE3), combined with a *treSA* from *P. putida* KT2440 in bicistronic construct | this study |
| pSEVA228-galP-TreF | Expression plasmid for *galP* from *E. coli* *E. coli* DH5α, combined with a *treSA* from *P. putida* KT2440 in bicistronic construct | this study |

As can be seen in Fig. 4B, low sugar concentrations were found for all transport systems, even for the negative control (wildtype TreB alone). This residual maltose/trehalose peak is probably a result of unspecific secretion, cell lysis or due to centrifugation of the cells. Apart from that, the expression plasmids pSEVA228-setA and pSEVA228-setA-treS showed an increased secretion of sugars, indicating that SetA can also transport trehalose. It is very likely that in the experiment with plasmid pSEVA228-setA-treS, maltose and trehalose were both secreted, but as maltose and trehalose show a similar retention time in our HPLC method, the disaccharides could not be separated in this experiment. TreB in combination with TreS also led to a secretion that was slightly higher than the negative control with the TreB wildtype enzyme. Facilitated transport of maltose through TreB has been described as a side reaction (Decker et al., 1999) and additionally, facilitated transport is not pushed by the proton motive force. This might explain the minor capability of TreB to transport maltose compared to SetA. Since trehalose is a more interesting and valuable sugar than maltose, we continued our cultivation studies with the strain *C. necator* H16 (pSEVA228-setA).

The finding that SetA can secrete trehalose also raised the question whether or not this protein is responsible for the secretion of trehalose in other bacteria that are naturally accumulating trehalose in the culture medium like *Corynebacterium glutamicium* (Wittmann et al., 2004) or various *Micrococcus strains* (Kizawa et al., 1995). In the two strains, *C. glutamicum* ATCC 13032 and *Micrococcus luteus* NCTC 2665, no significant homolog of SetA could be found. Therefore, the proteins involved in trehalose export remains elusive in these strains, but might have the potential to be a further source of a trehalose transport protein.

### Engineering of the GITE-motif of the PTS transport subunit for decoupling was not predictable

The lack of trehalose secretion by TreB variants shows that the effect of mutations is not predictable enough for rational engineering of the PTS transport subunits for facilitated transport. Interestingly, although the TreB(E385A) protein did not show any facilitated transport, the strain overexpressing this protein grew considerably slower (data not shown). If this approach is to be followed in the future, it would be interesting to perform semi-rational protein engineering of residues in and near the GITE-motif with saturation mutagenesis. Alternatively, a selection scheme for facilitated transport of TreB could be developed, for example by coupling the TreB gene with the reversible trehalose synthase TreS and a maltose phosphorylase and thereby allowing growth only with a facilitated transport of trehalose by TreB without phosphorylation. Such selection schemes have been successful in *E. coli* in other studies (Otte et al., 2003; Ruijter et al., 1992).

### Sugar/H^+^-symporters are not suitable for sugar secretion in bacteria with a normal PMF

In the experiments with constructs using MalY as the secretion protein, no significant amount of sugar have been detected as well. We tested the activity of TreS by enzyme assays with the lysate of these cells to exclude a problem with the conversion of trehalose to maltose. As in *C. necator*, it is expected that the proton motive force (PMF) at pH 7 promotes higher proton concentrations on the periplasmic site of the membrane, maltose secretion with a symporter seems not very likely. This leaves the question: Why did sugar transport work with the sucrose/H^+^-symporter CscB in *S. elongatus* in the first place? As Ducat *et al.* (Ducat et al., 2012) noted, the membrane potential is reversed in *S. elongatus* because of the high pH outside of the cell. However, this does also mean that ATP cannot be generated at the inner membrane by canonical proton-translocating ATP-Synthases. It is likely that ATP is rather produced at the thylakoid membrane of these organisms and that the plasma membrane does not have an active transmembrane gradient. This might also contribute to the high pH tolerance of this organism that can thrive at pH > 9 (Mortezaeikia et al., 2016) and is in line with the transport of carbonate via a Na^+^-gradient or in an ATP dependent manner (Schwarz et al., 2011). In any case, for a bacterium with a text-book proton motive force (PMF) like *C. necator* H16, an H^+^/symporter is not supposed to function as a secretion protein. SetA, in contrast, was found to act as an antiporter (Liu, Miller, Gosink, et al., 1999), and thus uses the PMF in the other direction, i.e. for export of sugar.

### Production of trehalose from organic substrates by engineered strains

After the initial screenings for trehalose secretion, we next characterized the *C. necator* (pSEVA228-setA) strain in shake flasks using different substrates. We were especially interested whether trehalose formation is a general feature in conditions of salt stress. Apart from fructose which will be metabolized by the Entner-Doudoroff pathway in *C. necator*, we also tested acetate and succinate as substrates that will cause *C. necator* to run gluconeogenesis. In addition, glycerol was tested as a cheap substrate that can be metabolized natively by *C. necator*, and even acts as a good substrate when the strain is engineered in this respect (Fukui et al., 2014). The results of shakeflasks experiments are shown in Fig. 5.

**Figure 5:**
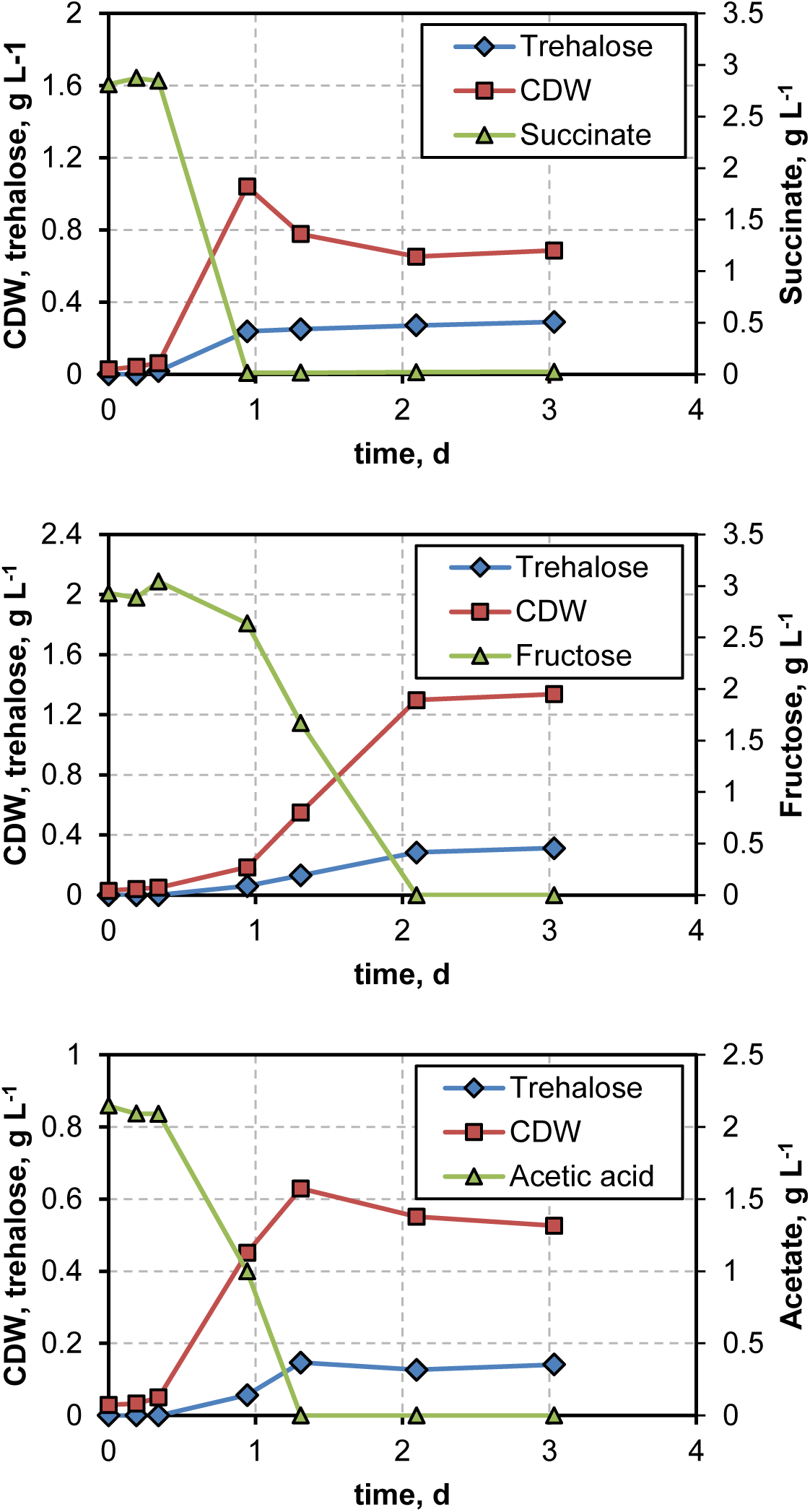
Growth and trehalose formation of *C. necator* H16 (pSEVA228-setA) over time using different substrates. Experiments have been run in shake flasks with 25 mL of MR medium at 30°C and 220 rpm. *setA* was expressed via the XylS/Pm system in plasmid pSEVA228 using 0.1 mM 3-MB at the time of inoculation.

These experiments clearly show that trehalose can also be produced and secreted using non-carbohydrate substrates. Trehalose production from carbohydrates is an already established process (Cai et al., 2018), so using alternative substrates, like glycerol might be an interesting alternative. Growth on glycerol of wildtype *C. necator* was extremely slow in our experiments as has also been previously reported by others (Fukui et al., 2014). After a very long lag-phase of more than 7 days, we only captured the culture after reaching stationary phase 15 days after inoculation. Nevertheless, the mere production of trehalose could be shown in this case from glycerol as well. Table 2 summarizes the concentrations of trehalose formed, the cell dry weight and the respective product/substrate yields for all the substrates after reaching the stationary phase. Fructose and succinate lead to higher biomass and trehalose yields than acetate, which makes sense when considering its metabolism: Using Acetyl-CoA synthetase, 2 equivalents of ATP are necessary for the conversion of acetate to Acetyl-CoA which in turn has to be processed via the glyoxylate-shunt to succinate. This means that directly using succinate as a substrate “costs” the cell 4 ATP less than the equivalent of acetate. Even when considering that one NADH is generated in the glyoxylate shunt, succinate will be the more energy efficient substrate per gram and also requires fewer enzymatic conversion steps and hence less protein investment. Fructose and glycerol are supposed to be the most attractive substrate for *C. necator* as both feed into glycolysis which generates ATP, in contrast to gluconeogenesis which is necessary for growth on acetate or succinate and consumes ATP.

The product/substrate yield *Y*_*P/S*_ of glycerol is probably slightly overestimated since there should have been some evaporation after 15 days that is difficult to measure. However, the value of 0.137 is in line with fructose that also enters the metabolism at a similar level (i.e. glycerol is directly converted into a C3-sugar). The product/substrate yields of around 0.1 g/g for fructose, succinate and glycerol are excellent starting values for further optimization: It has to be stressed that by the expression of *setA*, we only implemented a pull strategy - i.e. trehalose production is increased by draining the product from the cell - but did not yet push the flux in the direction of trehalose. Enforcing the flux in the direction of trehalose by overexpression of the trehalose synthesis genes, for example *otsA* and *otsB*, should increase the production rate and yields considerably. This will be part of future research projects in our group.

### Production of trehalose from a H_2_/CO_2_ mixture by engineered strains

Finally, we tested the production of trehalose with *C. necator* H16 (pSEVA228-setA) from gaseous substrates. Cultivations with a H_2_/CO_2_/O_2_ gas-mixture in modified MR-medium resulted in formation of up to around 0.47 g/L trehalose when using 150 mM NaCl as depicted in Fig. 6. No side product formation was observed. Lower salt concentrations resulted in lower trehalose production while higher salt concentrations led to higher trehalose and lower biomass formation. These data confirm the salt dependence of trehalose formation in this strain from gaseous substrates as well.

**Figure 6:**
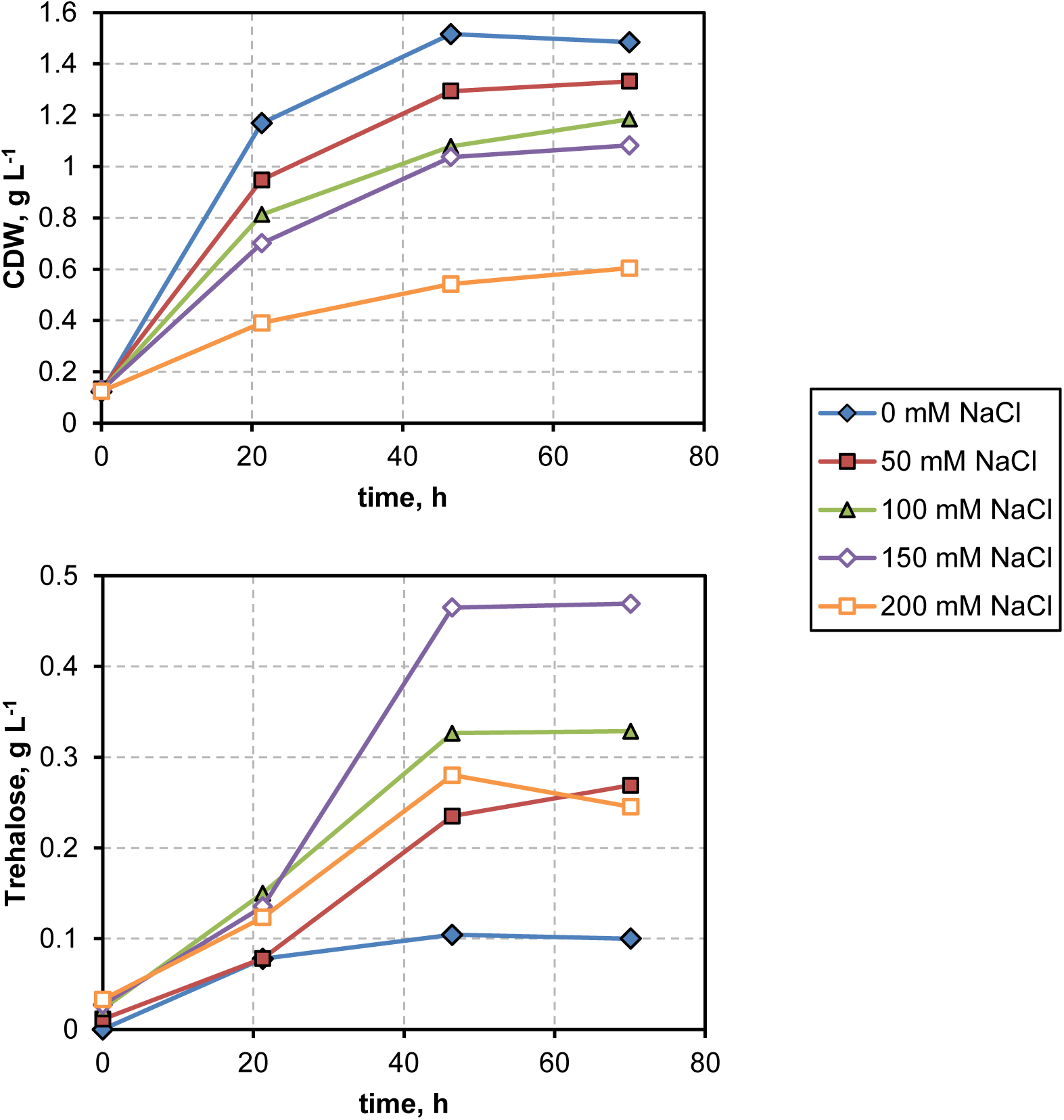
Trehalose and biomass concentration over time when culturing *C. necator* H16 (pSEVA228-setA) on a 6:1:1 mixture of H_2_:CO_2_:O_2_ in MR mineral medium. Varying amounts of salt were added (0-200 mM) and the cultures were induced with 0.1 mM 3-MB. Cultivations were performed in 500 mL screw-bottles with rubber sealings and 25 mL culture volume.

As it was not possible to measure residual hydrogen and leakage of hydrogen in the gas phase, it is difficult to estimate the yields of trehalose. Judging from the ratio of trehalose and biomass, the cultivations with the gas mixture as substrate formed 0.434 g trehalose per gram of biomass, which is in the same order of magnitude as for the heterotrophic substrates (0.233 g/g for acetate, 0.279 for succinate, 0.233 for fructose and 0.305 for glycerol). In fact, when fed on gaseous substrates, more carbon was directed to trehalose than in the heterotrophic cultivations. We observed, however, flocculation of biomass in the gas-based fermentations and thus the value might be over-estimated. Using the very simplified assumption that trehalose and biomass have similar yields from hydrogen and that a stoichiometry of 3.154 g(CDW)/mol(H_2_) can be reached as determined in former studies (Tanaka et al., 1995), the product/substrate yield can be estimated to be in the order of 0.953 g(Trehalose)/mol(H_2_). How does that compare to the maximal theoretical yield? If we assume that the Calvin cycle needs around 9-12 ATP to form one mol of Glyeraldehyde-3-phosphate (GAP) (Bar-Even et al., 2010) and ATP can be generated from hydrogen with a stoichiometry of 2.5 ATP/H_2_ (using the cytoplasmic hydrogenase of *C. necator* H16 and a ATP/NADH ratio of 2.5), we get a maximal yield of 1 mol trehalose per 26.8-31.6 mol H_2_ which equals to 10.8-12.8 g(Trehalose)/mol(H_2_). We can therefore estimate that 7.5-8.8% of the maximal theoretical yield was reached in our experiments. Even though we did not engineer enhanced trehalose formation, but only leakage from the cell, this is already a good starting point and we expect to come closer to the maximal values when further metabolic engineering is applied.

## Conclusion

In this study, we show the production of the sugar trehalose from carbon dioxide, hydrogen and oxygen gas by engineering the hydrogen-oxidizing bacterium *C. necator H16* for trehalose leakage for the first time. Trehalose is used by the food and pharma industry and is more valuable than, for example, glucose or sucrose (food grade trehalose sells for ∼3$/kg (L O’Brien-Nabors, 2016) in contrast to glucose and sucrose that sell below 1$/kg (Elvers, 2017)), but also than other products like ethanol or methanol (Straathof & Bampouli, 2017) that can be produced from synthesis gas and are already commercialized. Apart from that, the process is well suited for the production of ^13^C-labeled sugar since the price ^13^C-labeled CO_2_ is magnitudes lower per carbon and by using a surplus of hydrogen, a high yield of ^13^CO_2_ should be possible. Another interesting use-case are scenarios where hydrogen gas and electricity are available, but space is sparse like on space missions: edible sugar could be produced without the need of large areas for plant growth and with a possibly better efficiency.

While finalizing this manuscript for publication, we noticed a similar study had been published on bioRxiv (Nangle et al., 2020) in which, among other products, sucrose was produced from a gaseous mixture by expression of sucrose synthesis genes and a sucrose porin. Sugar produced from CO_2_ was successfully used to feed heterotrophic processes as shown by Nangle *et al.* and former studies (Ducat et al., 2012; Löwe et al., 2017), but this biotechnologically produced sugar (by *S. elongatus* or *C. necator*) will likely be too expensive in the beginning in a commercial setting. Therefore, engineering *C. necator* and other hydrogen-oxidizing bacteria directly will probably be the better route. *C. necator* is one of the best understood strains of the hydrogen-oxidizing bacteria and therefore well suited for the production of energy-demanding products such as trehalose. Compared to other hydrogen utilizing microorganisms like acetogens or methanogens, hydrogen-oxidizing bacteria can breathe oxygen and thereby generate the required ATP. Therefore, we expect a lot more processes for the production of ATP-demanding compounds in the future with this remarkable class of microorganisms.

## Material and Methods

### Bioinformatic analysis

In order to identify genes responsible for glycogen or sugar synthesis, a homology search was performed using the Basic Local Alignment Search Tool (BLAST) (Altschul et al., 1990). For each protein, a template was chosen from an organism in which the function of the respective protein was characterized in former studies: Since in *C. glutamicum* ATCC 13032, homologs to most genes could be found, the protein sequences of this organism were used as a reference except for the sequences GlgA and TreF that were not present in its genome, and for the GH15-family hydrolase that was not characterized. For these proteins, sequences from *E. coli* K-12 MG1655 (GlgA and TreF) or *C. necator* H16 (GH15-family hydrolase) were used as a template. Homology was evaluated based on the score that was normalized to the score of the template itself. The complete analysis including gene locus names and the outcomes of the homology search, can be found as a table in the supplement.

### Construction of expression plasmids

All expression plasmids used in this study derive from the XylS/Pm-based expression plasmid pSEVA228 of the Standard European Vector Architecture (SEVA) library (Martínez-García et al., 2015). In general, restriction/ligation cloning was used for the construction of plasmids. In brief: Genes of interest were amplified with PCR from genomic DNA (gDNA) templates. gDNA was extracted from *E. coli* DH5α, *E. coli* BL21(DE3), *P. putida* KT2440 or *N. lactamica* DSM 4691 using the “Genomic DNA from tissue” kit by MACHEREY-NAGEL GmbH & Co. KG (Germany) or was directly ordered from the DSMZ GmbH (Germany). Forward primers included a restriction enzyme recognition site, followed by the Ribosome Binding Site (RBS) AGGAGGCTTCAT, directly followed by the start codon. A list of all used primers and other oligonucleotides can be found in Table 3. PCR products and pSEVA228 plasmids were purified, digested with suitable restriction enzymes and ligated with T4 ligase.

For the site-specific mutations in the *treB* gene, overlap extension PCR (OE-PCR) was used in three rounds of PCR (Higuchi et al., 1988): In a first round, up- and downstream fragments (1.2 kB upstream, 0.3 kB downstream) were PCR-amplified using specific primers carrying the site-specific base changes. After gel purification, equal molarities of both fragments were mixed in a new Q5 PCR reaction without primers, using an annealing temperature of 60°C and an elongation time corresponding to the full length constructs for 15 cycles. 8 µL of this reaction was directly used without purification in the third and final round of PCR in a 50 µL reaction including the flanking primers treB_E385A_fw2 and treB_V383N_fw2 with restriction enzyme recognition sites. This was restriction digested, purified and ligated as described above.

All purification and extraction steps were performed using kits by MACHEREY-NAGEL or NIPPON Genetics EUROPE GmbH (Germany). All enzymes were ordered from New England Biolabs GmbH (Germany). A complete list of plasmids used in this study can be found in Table 4.

### Transformation & conjugation

Ligated DNA was transformed into competent *E. coli* DH5α λ-*pir* following the method of Chung *et al.* (Chung et al., 1989). Resulting colonies were checked with colony-PCR, followed by sequencing using suitable primers. For expression in *C. necator* H16, the plasmids were then transferred via conjugation with the sitting drop method using the helper strain *E. coli* HB101 (pRK600) as described by others (de Lorenzo & Timmis, 1994; Löwe et al., 2018) into *C. necator H16. C. necator* H16 strains with the desired plasmids were then selected by streaking the conjugation mixture on either minimal medium with 300 mg/L kanamycin using citrate as a carbon source or in LB medium with 300 mg/L kanamycin and 100 mg/L streptomycin.

### Cultivation media and heterotrophic cultivation procedures

*E. coli* DH5α and BL21(DE3) and *C. necator* H16 cultures for plasmid and genomic DNA extraction, as well as for transformation, were grown in Lysogeny-Broth (LB) medium. *C. necator* was additionally cultured in minimal medium: For the initial screenings of the different transport proteins, a modified low-phosphate minimal medium was used (de Aragão et al., 1996) as trehalose and phosphate share a similar retention time in our HPLC method and peaks are thus likely to overlap making quantification difficult at high phosphate concentrations. The medium consisted of (per liter): 0.19 g EDTA, 0.06 g ferrous ammonium citrate, 0.5 g MgSO_4_, 0.01 g of CaCI_2_·2H_2_O and 5.0 g of (NH_4_)_2_SO_4_. Additionally, 1 mL/L of a trace element solution was added, consisting of (per liter): 0.3 g of H_3_BO_3_, 0.2 g of CoCl_2_·6H_2_O, 0.1 g of ZnSO_4_·7H_2_O, 0.03 g of MnCI_2_·4H_2_O, 0.03 g of Na_2_MoO_4_·2H_2_O, 0.02 g of NiCI_2_·6H_2_O and 0.01 g of CuSO_4_·5H_2_O. For the heterotrophic and autotrophic cultivations of *C. necator* H16 (pSEVA228-setA), a phosphate-rich, modified MR medium (Y. Lee & Lee, 1996) was used to buffer the acidic carbon source and the carbonate in autotrophic cultivations. Modified MR medium consisted of (per liter): 13.5 g KH_2_PO_4_, 4.0 g (NH_4_)_2_HPO_4_, 1.4 g MgSO_4_·7H_2_O. Citric acid was replaced with the carbon source of interest, e.g. sodium acetate, fructose etc. For the sake of simplification, the same trace element solution was used as in the low-phosphate medium. The missing iron solution was supplied as a 100x stock solution of (per liter) 19 g EDTA, 6 g ferrous ammonium citrate.

Cultures in shake flasks were incubated at 37°C for *E. coli* and 30°C for *C. necator* and *P. putida* in a rotary shaker (ThermoFisher, USA) at a rotation frequency of 220 rpm.

Antibiotics were used when necessary in solid or liquid media: 300 mg/L kanamycin for *C. necator* (50 mg/L for *E. coli*), 34 mg/L chloramphenicol (*E. coli*), 100 mg/L streptomycin (*E. coli).* The inducer 3-methylbenzoic acid was in general used at a concentration of 0.1 mM.

### Autotrophic cultivation of *C. necator* H16 (pSEVA228-setA)

Cultivation of *C. necator* H16 (pSEVA228-setA) were carried out in air-tight butyl-rubber sealed 500 mL security-coated glass bottles with a working volume of 25 mL. Bottles and sealings were autoclaved separately. After sterilely filling medium into the bottles, they were tightly sealed and gassed with a mixture of 6:1 of H_2_:CO_2_ for 5 min through a syringe needle. A second syringe needle was stung through the rubber to release the pressure. After that, 71 mL of pure oxygen were added with a sterile syringe. Inoculation and sampling were also achieved through the rubber sealing with sterile syringes. Each bottle was inoculated with 0.4 mL of LB-grown *C. necator* pre-cultures. Bottles were shaken at a temperature of 30°C and a frequency of 100 rpm in a shaking incubator. All critical operations were done in an explosion protected fume hood.

### Optical density and sugar analytics

Optical densities (OD) were measured at 600 nm in a microplate reader (TECAN, Switzerland) using a volume of 200 µL in transparent microplates. ODs were correlated to optical densities measured in a photometer with a light path of 1 cm and then to CDW using a correlation described by others (Grunwald et al., 2015).

Sugar concentrations were measured following a previously published protocol (Löwe et al., 2018) with a Shodex SH1011 column. In brief, an Agilent 1100 series (Waldbronn, Germany) system was used with filtered 0.5 mM H_2_SO_4_ as mobile phase, a flow rate of 0.45 mL/min and a temperature of 30°C. 20 µL of filtered and diluted sample were injected and analyzed by their refractive index (RI) signal. In cultivation experiments with high-phosphate medium, trehalose was hydrolyzed prior to analysis with the cell-lysates containing the trehalase TreF from *E. coli*. The cell lysate was produced as follows: *E. coli* (pSEVA228-galP-TreF) was grown in 25 mL of LB with 50 mg/L kanamycin and 0.1 mM of the inducer 3-MB. When cultures entered stationary phase, the fermentation broths were centrifuged and up-concentrated to 1 mL in phosphate buffered saline (PBS). Cells were then mixed with glass beads and lysed in a ball mill (RETSCH GmbH, Germany) for 15 min. Cell lysates were centrifuged and the supernatant was stored at 8°C. For the hydrolysis of trehalose, mixtures with 10% lysate and 90% sample were incubated for 2 hours, filtered and analyzed by HPLC.

## Supporting information

Details of the bioinformatic analysis

## Acknowledgments

We would like to thank Prof. Dirk Weuster-Botz for the opportunity to use the infrastructure for gas fermentation at the institute of Biochemical Engineering at the Technical University of Munich.

## Author contributions

H.L. wrote the manuscript. H.L. designed the experiments and performed all experimental work with support by M.B. The work was supervised by K.P.G and A.K. who were also involved in editing of the manuscript.

## Notes

### Competing Interest Statement

The authors have declared no competing interest.

## Bibliography

Altschul, S. F., Gish, W., Miller, W., Myers, E. W., & Lipman, D. J. (1990). Basic local alignment search tool. Journal of Molecular Biology, 215(3), 403–410. https://doi.org/ https://doi.org/10.1016/S0022-2836(05)80360-2

Bar-Even, A., Noor, E., Lewis, N. E., & Milo, R. (2010). Design and analysis of synthetic carbon fixation pathways. Proceedings of the National Academy of Sciences of the United States of America, 107(19), 8889–8894. https://doi.org/10.1073/pnas.0907176107

Belda, E., van Heck, R. G. a, Lopez-Sanchez, M. J., Cruveiller, S., Barbe, V., Fraser, C., Klenk, H.-P., Petersen, J., Morgat, A., Nikel, P. I., Vallenet, D., Rouy, Z., Sekowska, A., Martins Dos Santos, V. a P., de Lorenzo, V., Danchin, A., & Médigue, C. (2016). The revisited genome of *Pseudomonas putida* KT2440 enlightens its value as a robust metabolic chassis. Environmental Microbiology, 00, 1–22. https://doi.org/10.1111/1462-2920.13230

Burg, M. B., & Ferraris, J. D. (2008). Intracellular organic osmolytes: Function and regulation. Journal of Biological Chemistry, 283(12), 7309–7313. https://doi.org/10.1074/jbc.R700042200

Cai, X., Seitl, I., Mu, W., Zhang, T., Stressler, T., Fischer, L., & Jiang, B. (2018). Biotechnical production of trehalose through the trehalose synthase pathway: current status and future prospects. Applied Microbiology and Biotechnology, 102(7), 2965–2976. https://doi.org/10.1007/s00253-018-8814-y

Carroll, J. D., Pastuszak, I., Edavana, V. K., Pan, Y. T., & Elbein, A. D. (2007). A novel trehalase from Mycobacterium smegmatis - Purification, properties, requirements. FEBS Journal, 274(7), 1701–1714. https://doi.org/10.1111/j.1742-4658.2007.05715.x

Chung, C. T., Niemela, S. L., & Miller, R. H. (1989). One-step preparation of competent Escherichia coli: transformation and storage of bacterial cells in the same solution. Proceedings of the National Academy of Sciences of the United States of America, 86(7), 2172–2175. http://www.ncbi.nlm.nih.gov/pmc/articles/PMC286873/

Claassens, N. J., Cotton, C. A. R., Kopljar, D., & Bar-Even, A. (2019). Making quantitative sense of electromicrobial production. Nature Catalysis, 2(5), 437–447. https://doi.org/10.1038/s41929-019-0272-0

Clausen, L. R., Houbak, N., & Elmegaard, B. (2010). Technoeconomic analysis of a methanol plant based on gasification of biomass and electrolysis of water. Energy, 35(5), 2338–2347. https://doi.org/10.1016/j.energy.2010.02.034

de Aragão, G. M. F., Lindley, N. D., Uribelarrea, J. L., & Pareilleux, A. (1996). Maintaining a controlled residual growth capacity increases the production of PHA copolymers by *Alcaligenes eutrophus*. Biotechnology Letters, 18(8), 937–942.

de Lorenzo, V., & Timmis, K. N. (1994). Analysis and construction of stable phenotypes in gram-negative bacteria with Tn5- and Tn10-derived minitransposons. Methods in Enzymology, 235, 386—405. https://doi.org/10.1016/0076-6879(94)35157-0

Decker, K., Gerhardt, F., & Boos, W. (1999). The role of the trehalose system in regulating the maltose regulon of Escherichia coli. Molecular Microbiology, 32(4), 777–788. https://doi.org/10.1046/j.1365-2958.1999.01395.x

Derkaoui, M., Antunes, A., Nait Abdallah, J., Poncet, S., Mazé, A., Pham, Q. M. M., Mokhtari, A., Deghmane, A. E., Joyet, P., Taha, M. K., & Deutscher, J. (2016). Transport and catabolism of carbohydrates by neisseria meningitidis. Journal of Molecular Microbiology and Biotechnology, 26(5), 320–332. https://doi.org/10.1159/000447093

Ducat, D. C., Avelar-Rivas, J. A., Way, J. C., & Silver, P. a. (2012). Rerouting carbon flux to enhance photosynthetic productivity. Applied and Environmental Microbiology, 78(8), 2660–2668. https://doi.org/10.1128/AEM.07901-11

Elvers, D. B. (2017). Ullmann’s Food and Feed, Volume 2 (D. B. Elvers (ed.)). John Wiley & Sons. https://books.google.de/books?id=Xwe3DQAAQBAJ

European Commission. (2020). Sustainable Europe Investment Plan, European Green Deal Investment Plan, COM(2020) 21 final. https://doi.org/10.1017/CBO9781107415324.004

Fukui, T., Mukoyama, M., Orita, I., & Nakamura, S. (2014). Enhancement of glycerol utilization ability of Ralstonia eutropha H16 for production of polyhydroxyalkanoates. Applied Microbiology and Biotechnology, 98(17), 7559–7568. https://doi.org/10.1007/s00253-014-5831-3

Grunwald, S., Mottet, A., Grousseau, E., Plassmeier, J. K., Popović, M. K., Uribelarrea, J. L., Gorret, N., Guillouet, S. E., & Sinskey, A. (2015). Kinetic and stoichiometric characterization of organoautotrophic growth of Ralstonia eutropha on formic acid in fed-batch and continuous cultures. Microbial Biotechnology, 8(1), 155–163. https://doi.org/10.1111/1751-7915.12149

He, Y., Wang, K., & Yan, N. (2014). The recombinant expression systems for structure determination of eukaryotic membrane proteins. Protein & Cell, 5(9), 658–672. https://doi.org/10.1007/s13238-014-0086-4

Higuchi, R., Krummel, B., & Saiki, R. (1988). A general method of in vitro preparation and specific mutagenesis of dna fragments: Study of protein and DNA interactions. Nucleic Acids Research, 16(15), 7351–7367. https://doi.org/10.1093/nar/16.15.7351

Joseph, T. C., Rajan, L. A., Thampuran, N., & James, R. (2010). Functional characterization of trehalose biosynthesis genes from E. coli: An osmolyte involved in stress tolerance. Molecular Biotechnology, 46(1), 20–25. https://doi.org/10.1007/s12033-010-9259-4

Kanamori, Y., Saito, A., Hagiwara-Komoda, Y., Tanaka, D., Mitsumasu, K., Kikuta, S., Watanabe, M., Cornette, R., Kikawada, T., & Okuda, T. (2010). The trehalose transporter 1 gene sequence is conserved in insects and encodes proteins with different kinetic properties involved in trehalose import into peripheral tissues. Insect Biochemistry and Molecular Biology, 40(1), 30–37. https://doi.org/10.1016/j.ibmb.2009.12.006

Kessler, B., Lorenzo, V. de, & Timmis, K. (1992). A general system to integratelacZ fusions into the chromosomes of gram-negative eubacteria: regulation of the Pm promoter of theTOL plasmid studied with all. Molecular and General Genetics …, 233(1), 293–301. http://link.springer.com/article/10.1007/BF00587591

Kizawa, H., Miyazaki, J. ichi, Yokota, A., Kanegae, Y., Miyagawa, K. ichiro, & Sugiyama, Y. (1995). Trehalose production by a strain of micrococcus varians. Bioscience, Biotechnology and Biochemistry, 59(8), 1522–1527. https://doi.org/10.1271/bbb.59.1522

Lee, S., Ju, H. K., Machunda, R., Uhm, S., Lee, J. K., Lee, H. J., & Lee, J. (2015). Sustainable production of formic acid by electrolytic reduction of gaseous carbon dioxide. Journal of Materials Chemistry A, 3(6), 3029–3034. https://doi.org/10.1039/c4ta03893b

Lee, Y., & Lee, S. Y. (1996). Enhanced production of poly(3-hydroxybutyrate) by filamentation-suppressed recombinantEscherichia coli in a defined medium. Journal of Environmental Polymer Degradation, 4(2), 131–134. https://doi.org/10.1007/BF02074874

Lin, P. C., Zhang, F., & Pakrasi, H. B. (2020). Enhanced production of sucrose in the fast-growing cyanobacterium Synechococcus elongatus UTEX 2973. Scientific Reports, 10(1), 1–8. https://doi.org/10.1038/s41598-019-57319-5

Liu, J. Y., Miller, P. F., Gosink, M., & Olson, E. R. (1999). The identification of a new family of sugar efflux pumps in Escherichia coli. Molecular Microbiology, 31(6), 1845–1851. https://doi.org/10.1046/j.1365-2958.1999.01321.x

Liu, J. Y., Miller, P. F., Willard, J., & Olson, E. R. (1999). Functional and biochemical characterization of Escherichia coli sugar efflux transporters. Journal of Biological Chemistry, 274(33), 22977–22984. https://doi.org/10.1074/jbc.274.33.22977

Löwe, H., Hobmeier, K., Moos, M., Kremling, A., & Pflüger-Grau, K. (2017). Photoautotrophic production of polyhydroxyalkanoates in a synthetic mixed culture of *Synechococcus elongatus* cscB and *Pseudomonas putida* cscAB. Biotechnology for Biofuels, 10(1). https://doi.org/10.1186/s13068-017-0875-0

Löwe, H., Sinner, P., Kremling, A., & Pflüger-Grau, K. (2018). Engineering sucrose metabolism in Pseudomonas putida highlights the importance of porins. Microbial Biotechnology. https://doi.org/10.1111/1751-7915.13283

Martínez-García, E., Aparicio, T., Goñi-Moreno, A., Fraile, S., & de Lorenzo, V. (2015). SEVA 2.0: an update of the Standard European Vector Architecture for de-/re-construction of bacterial functionalities. Nucleic Acids Research, 43(Database issue), D1183–9. https://doi.org/10.1093/nar/gku1114

Mortezaeikia, V., Yegani, R., & Tavakoli, O. (2016). Membrane-sparger vs. membrane contactor as a photobioreactors for carbon dioxide biofixation of Synechococcus elongatus in batch and semi-continuous mode. Journal of CO2 Utilization, 16, 23–31. https://doi.org/10.1016/j.jcou.2016.05.009

Nangle, S. N., Ziesack, M., Buckley, S., Trivedi, D., Loh, D. M., Nocera, D. G., & Silver, P. A. (2020). Valorization of CO2 through lithoautotrophic production of sustainable chemicals in *Cupriavidus necator*. BioRxiv, 2020.02.08.940007. https://doi.org/10.1101/2020.02.08.940007

O’Brien-Nabors, L. (2016). Alternative Sweeteners (Lyn O’Brien-Nabors (ed.); 4th, illus ed.). CRC Press. https://books.google.de/books?id=5yHOBQAAQBAJ

Otte, S., Scholle, A., Turgut, S., & Lengeler, J. W. (2003). Mutations Which Uncouple Transport and Phosphorylation in the. Microbiology, 185(7), 2267–2276. https://doi.org/10.1128/JB.185.7.2267

Peng, Y., Kumar, S., Hernandez, R. L., Jones, S. E., Cadle, K. M., Smith, K. P., & Varela, M. F. (2009). Evidence for the transport of maltose by the sucrose permease, CscB, of escherichia coli. Journal of Membrane Biology, 228(2), 79–88. https://doi.org/10.1007/s00232-009-9161-9

Rothschild, A., & Dotan, H. (2017). Beating the Efficiency of Photovoltaics-Powered Electrolysis with Tandem Cell Photoelectrolysis. ACS Energy Letters, 2(1), 45–51. https://doi.org/10.1021/acsenergylett.6b00610

Ruhal, R., Kataria, R., & Choudhury, B. (2013). Trends in bacterial trehalose metabolism and significant nodes of metabolic pathway in the direction of trehalose accumulation. Microbial Biotechnology, 6(5), 493–502. https://doi.org/10.1111/1751-7915.12029

Ruijter, G. J. G., Van Meurs, G., Verwey, M. A., Postma, P. W., & Van Dam, K. (1992). Analysis of mutations that uncouple transport from phosphorylation in enzyme II(Glc) of the Escherichia coli phosphoenolpyruvate-dependent phosphotransferase system. Journal of Bacteriology, 174(9), 2843–2850. https://doi.org/10.1128/jb.174.9.2843-2850.1992

Sahintoth, M., Frillingos, S., Lengeler, J. W., & Kaback, H. R. (1995). Active Transport by the CscB Permease in Escherichia coli K-12. Biochemical and Biophysical Research Communications, 208(3), 1116–1123.

Schlegel, S., Klepsch, M., Gialama, D., Wickström, D., Slotboom, D. J., & De Gier, J. W. (2010). Revolutionizing membrane protein overexpression in bacteria. Microbial Biotechnology, 3(4), 403–411. https://doi.org/10.1111/j.1751-7915.2009.00148.x

Schwarz, D., Nodop, A., Hüge, J., Purfürst, S., Forchhammer, K., Michel, K. P., Bauwe, H., Kopka, J., & Hagemann, M. (2011). Metabolic and transcriptomic phenotyping of inorganic carbon acclimation in the cyanobacterium synechococcus elongatus PCC 7942. Plant Physiology, 155(4), 1640–1655. https://doi.org/10.1104/pp.110.170225

Straathof, A. J. J., & Bampouli, A. (2017). Potential of commodity chemicals to become bio-based according to maximum yields and petrochemical prices. Biofuels, Bioproducts and Biorefining, 11, 798–810. https://doi.org/10.1002/bbb

Sun, X., Atiyeh, H. K., Huhnke, R. L., & Tanner, R. S. (2019). Syngas fermentation process development for production of biofuels and chemicals: A review. Bioresource Technology Reports, 7(July). https://doi.org/10.1016/j.biteb.2019.100279

Tanaka, K., Ishizaki, A., Kanamaru, T., & Kawano, T. (1995). Production of poly(D-3-hydroxybutyrate) from CO(2), H(2), and O(2) by high cell density autotrophic cultivation of *Alcaligenes eutrophus*. Biotechnology and Bioengineering, 45, 268–275.

Thiel, K., Patrikainen, P., Nagy, C., Fitzpatrick, D., Pope, N., Aro, E. M., & Kallio, P. (2019). Redirecting photosynthetic electron flux in the cyanobacterium Synechocystis sp. PCC 6803 by the deletion of flavodiiron protein Flv3. Microbial Cell Factories, 18(1), 1–16. https://doi.org/10.1186/s12934-019-1238-2

Wittmann, C., Kiefer, P., & Zelder, O. (2004). Production with Sucrose as Carbon Source. Society, 70(12), 7277–7287. https://doi.org/10.1128/AEM.70.12.7277

Wolf, A., Krämer, R., & Morbach, S. (2003). Three pathways for trehalose metabolism in Corynebacterium glutamicum ATCC13032 and their significance in response to osmotic stress. Molecular Microbiology, 49(4), 1119–1134. https://doi.org/10.1046/j.1365-2958.2003.03625.x

Woodhouse, P. (2010). Beyond industrial agriculture? Some questions about farm size, productivity and sustainability. Journal of Agrarian Change, 10(3), 437–453. https://doi.org/10.1111/j.1471-0366.2010.00278.x

Yishai, O., Lindner, S. N., Gonzalez de la Cruz, J., Tenenboim, H., & Bar-Even, A. (2016). The formate bioeconomy. Current Opinion in Chemical Biology, 35, 1–9. https://doi.org/10.1016/j.cbpa.2016.07.005

Zhu, X. G., Long, S. P., & Ort, D. R. (2008). What is the maximum efficiency with which photosynthesis can convert solar energy into biomass? Current Opinion in Biotechnology, 19(2), 153–159. https://doi.org/10.1016/j.copbio.2008.02.004

